# Urban *Lasius niger* ants more readily accept low concentration sucrose solution than rural ants

**DOI:** 10.1101/2025.08.18.670873

**Authors:** Stanislav Stukalyuk, Mykola Kozyr, Yaroslav Sydorenko, Gema Trigos-Peral, Tomer J. Czaczkes

## Abstract

Urbanisation causes broad changes in biotic and abiotic factors, which are often detrimental to animals. While extensive attention has been focussed on food resources for pollinators, comparatively little research has examined resources for aphidophagous insects. Many ants, such as *Lasius niger*, are highly aphidophagous, relying on honeydew secretions for the majority of their carbohydrate intake. Ants will also often reject sugar solution which is of lower concentration than that they were expecting. Here, we ask whether *Lasius niger* ants from urban environments show differential acceptance to various sucrose solution molarities. We offered outgoing ants on active foraging trails drops of sucrose solutions over a range of molarities (0.125, 0.25, 0.5, 1, and 2M), and quantified acceptance. Acceptance was scored using a simple behavioural measure of whether the ant remained in contact with the sucrose drop for over 3 seconds (full acceptance), broke away from the drop within 3 seconds but remained in the area and eventually drank to satiation (partial acceptance), or did not drink to satiation and walked away (rejection). Urban ants showed a significantly higher acceptance of all sucrose molarities than rural ants, with this difference especially pronounced at lower molarities. This may indicate that they are under nutritional stress for carbohydrates. Ant responses to sucrose solutions may allow a straightforward measure of the availability of carbohydrate resources in the environment.

## Introduction

Urbanisation and land-use alterations are one of the main causes of the current global biodiversity crisis (Almond et al. 2022). Although the threats to biodiversity are diverse (e.g., ocean alteration, terrestrial land use, insecticides, invasive species; (Hald-Mortensen 2023), these threats largely result from environmental changes caused by human activities. Among these, land use change represents one of the most important drivers of biodiversity loss (Oliver and Morecroft 2014), primarily due to the combined effects of multiple abiotic and biotic pressures, such as habitat loss, fragmentation (Zhang et al. 2024), and the introduction of invasive species (Mollot et al. 2017).

Urbanization represents one of the most extreme forms of land-use transformation, although its full impact on native biodiversity remains insufficiently understood. As a result, urban ecological research has gained increasing relevance in recent years, driven by the urgent need to understand the functioning of these artificial ecosystems and to inform conservation strategies aimed at mitigating their widespread ecological consequences (McIntyre 2000). Urbanisation brings about profound physical and environmental changes—for example, the conversion of natural meadows into impermeable surfaces. Less visible, yet equally important, are alterations in soil chemistry, including elevated levels of heavy metals and other toxic compounds (Swain 2024). Urbanization also induces shifts in environmental conditions, such as increased exposure to artificial light at night (Hopkins et al. 2018) and the urban heat island effect (MacLachlan et al. 2017). Together, these changes have triggered a wide range of responses within urban biotic communities, spanning from shifts in community composition and interactions in biological networks (e.g. (Miles et al. 2019; Patterson et al. 2023; Marcacci et al. 2023; Stukalyuk and Maák 2023; Trigos-Peral and Reyes-López 2025) to behavioural and physiological adaptations at the individual level (e.g.(Ditchkoff et al. 2006; Meillère et al. 2015; Brans et al. 2018; Minias 2023; Tougeron and Sanders 2023). For instance, urbanization has been linked to declines in epiphyte diversity (Becker et al. 2017), reduced the health and performance of bumble bee colonies (Theodorou et al. 2022), intensified the offspring defence in urban populations of the Black Kite *Milvus migrans* (Kumar et al. 2018), reduced attraction to light in moths and shyness in spiders (Altermatt and Ebert 2016; Czaczkes et al. 2018) and alterations in animals phenology like diurnal spiders switching to hunt during the night facilitated by the artificial illumination in cities (Frank 2009) or changes in ant foraging activity and colony development (Trigos-Peral et al. 2024a). While some organisms have the capacity to adjust their behaviour and physiology in response to urban environmental conditions (Luniak 2004; Czechowski et al. 2012; McDonnell and Hahs 2015), enabling them to persist in these novel ecosystems, others – like those with low ecological plasticity – have faced more severe consequences, including the local extinction of their populations (Ancillotto et al. 2025, Dri et al. 2021, Duncan et al. 2011).

As previously mentioned, organisms respond to environmental changes in diverse ways, both in form and timescale—ranging from rapid behavioural adjustments that help overcome temporary conditions to slower evolutionary adaptations that persist across generations. Beyond the crucial role that both types of responses play in organisms survival, rapid behavioural responses can also offer practical applications with potential benefits for humans, namely bioindication. The use of bivalve molluscs response to decreases in water quality as an effective biomonitoring method in water quality monitoring systems globally is a widely cited example of a bioindicator (Vereycken and Aldridge 2023). However, aquatic organisms are not the only group capable of serving this function; soil organisms can also provide valuable insights into habitat status and quality (Viana et al. 2010; Morelli et al. 2021). Shifts in the functional traits of their populations often reflect changes in environmental conditions, owing to the strong association between species’ ecological requirements and habitat characteristics. This is particularly significant for organisms with limited dispersal capacity, which tend to be highly specialized to local conditions and, consequently, more sensitive to environmental changes—making them effective biological indicators (McGeoch 1998).

Although all organisms respond to changes in environmental conditions, those considered suitable as bioindicators must meet a specific set of criteria. They should be widely distributed, highly abundant, and relatively easy to sample. In addition, they should play a key functional role in ecosystems, exhibit high sensitivity to environmental changes, and display consistent responses that can be reliably interpreted (McGeoch 1998; Andersen 1999). Through the categorization in functional groups (Andersen 1995; Susilo et al. 2009) or directly using the specific ecology of the species, bioindicators provide a valuable information about the characteristics and conservation status of the habitats. For example, the occurrence of desert ants indicates a xerothermic habitat (de Pletincx et al. 2024), the presence of odonatans indicates presence of water bodies nearby (Carchini et al. 2007), the existence of ferns in an area is an indication of high moisture and mild temperatures (Della 2022), and different species of leaf litter fungi are associated to specific succession stage of woodlands (Prakash et al. 2015). Indeed, bioindicators have been used widely in ecological studies to evaluate changes in the environmental characteristics of the ecosystems or even the presence of pollutants (Viana et al. 2010; Ghannem et al. 2024).

Among all bioindicator organisms, ants are widely regarded as one of the most valuable and reliable. Their sensitivity to environmental disturbances, combined with their ease of sampling and taxonomic resolution, makes them particularly effective indicators of habitat quality and ecosystem change (Underwood and Fisher 2006). In broad terms, ant species are traditionally classified into two main bioindicator groups based on their niche preference and tolerance to varying environmental conditions: generalists, which exhibit broad ecological requirements and plasticity, and specialists, which have narrow ecological niches (Andersen 1999; Roig and Espadaler 2010; Vasconcelos et al. 2018; Trigos-Peral et al. 2020). While this classification is effective for assessing abiotic characteristics of habitats, it proves less efficient in detecting biotic changes, such as shifts in the availability and quality of nutritional resources.

The availability of abundant, high-quality food is crucial for organisms with limited nest mobility, such as ants. A scarcity of essential nutrients within their nesting niche can significantly impact ant populations, as a nutrient-rich diet is vital for colony development and the overall physiological status of ants and colonies (Csata and Dussutour 2019). For aphidicolous ant species like *Lasius niger*, aphid honeydew constitutes a key component of their diet. This sugary secretion offers a complex mixture of micronutrients important to the ants (Shik et al. 2014). As a result, ants actively select the most nutritious honeydew available to them. However, food source selection is not determined solely by nutritional content; previous feeding experience also plays a role (Wendt and Czaczkes 2017; Wendt et al. 2019).:Ants accustomed to richer food sources may exhibit reduced acceptance of alternatives with comparatively lower nutritional quality, as result of a negative incentive contrasts (Flaherty 1996). As such, the willingness of aphidicolous to accept low molarity sucrose concentrations should be a direct function of the sugar concentration or general quality of the honeydew they have been collecting, and this in turn may reflect the productivity – and thus stress levels – of the plant the aphids are feeding on.

This study aims to investigate potential shifts in aphid honeydew quality between urban and rural habitats by using the aphidicolous ant *Lasius niger* as a model species. Specifically, we seek to assess whether *L. niger* can serve as a bioindicator of the nutritional quality of available food sources by examining differences in dietary preferences by comparing the acceptance of sucrose solutions of varying concentrations between ant populations originating from highly urbanisation or low urbanisation environments.

## Methods

### Species and Localities

To assess whether ants can serve as reliable bioindicators of food quality in human-modified habitats, we used the generalist Holarctic ant species *Lasius niger* as model species. This aphidicolous species relies on honeydew produced by aphids as its main source of carbohydrates (Devigne and Detrain 2005; Novgorodova 2015). *L. niger* employs mass recruitment to food sources, so workers form stable foraging trails from the nest entrance to the exploited source (Devigne and Detrain 2005; Novgorodova 2015) and foraging trails can reach up to 2.9 m long (Stukalyuk & Akhmedov 2022), making them particularly easy to find. Moreover, the ecological plasticity of *Lasius niger* enables them to colonize a wide range of habitat types (Seifert 2007; Czechowski et al. 2012; Radchenko 2016), including highly modified habitats such as urban areas (Stukalyuk and Maák 2023). In such urban habitats, the species exhibits a marked dominance (Trigos-Peral et al. 2020), representing nearly 50% of the observed ant workers (Stukalyuk et al. 2020).

The study was performed in two habitat types differing in the degree of urbanization: urban and rural areas. Urban areas were characterized by green spaces embedded within the cityscape, including courtyards with paved roads, tiled pavements, and lawns with presence of a few trees; whereas rural areas were defined as areas of pastures, herbs, or spots surrounded by orchards. Both biomes were sampled across five different cities within the Kyiv district of Ukraine: Bucha, Boyarka, Irpin, Gostomel, and Vishneve. The GPS coodinates of all sampled locations can be found in the raw data (S1)

### Procedure

An active *Lasius niger* foraging trail was located by searching tree trunks for ants shuttling back and forth. Species identity was confirmed by examining samples of ants from each foraging trail under a stereo microscope, using identification keys in Radchenko (2016) and Seifert (2007). To perform the behavioural assay, a small acetate sheet (c. 10×10mm) was placed directly onto the trail, and a drop of sucrose solution was placed on the sheet. We tested the following sucrose molarities: 0.125, 0.25, 0.5, 1, and 2M, with a randomised presentation order. Only one drop of one solution was presented at a time. After 20 ants encountered the solution, the acetate sheet was removed and a new sucrose solution drop of a different molarity was presented. The experimenter was blind to solution type, although high molarity (1M, 2M) sucrose is viscous and thus can be differentiated from the other molarities even when unlabelled. The response of outgoing ants (heading towards the tree) when encountering the solution was noted. We recorded one of three responses, following Wendt et al. (2019):

- Full acceptance (scored as 1): The ant maintains its mandibles in contact with the sucrose drop for 3 seconds upon first contact with the drop.
- Partial acceptance (scored as 0.5): The ant breaks contact with the drop within 3 seconds of initial contact, but remains in the vicinity (c. 2cm radius) of the drop, returns to it frequently, or eventually drinks to satiation.
- Rejection (scored as 0): The ant breaks contact with the drop within 3 seconds, and continues on its outwards journey.

Solutions (all concentrations) were offered in patterned manner, so that all solutions were tested at each testing bout. All experiments were conducted in the shade, using a parasol to provide shade if necessary, and all data were collected between 9:00 and 12:00, between the 14^th^ and 27^th^ August 2024. In addition, the ambient air temperature, ambient humidity, tree trunk diameter, and ant trail traffic was measured for each colony. Ant traffic was measured by counting all ants passing by a point on the trail for 2 minutes.

We studied 9 urban colonies and 9 rural colonies from each municipality, testing 20 ants from each colony on each molarity, resulting in a total of 9000 ants tested.

### Statistical analysis

All code used in the analysis, and the code output, is provided in online supplement S1.

We analysed the data using Linear Mixed-Effect Models, with the glmmTMB package (Magnusson et al. 2020) in R (R Core Team 2023) via Rstudio (RStudio Team 2015). Acceptance responses were predicted by an interaction of food concentration and biome, with the addition of temperature and humidity as independent variables. Colony ID was added as a random effect. Our main analysis model formula was:

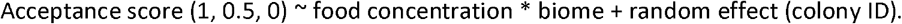

Differences in sucrose acceptance between habitat types were tested using a binomial model.

In addition, we explored the effect of environmental factors (humidity and temperature) and tree diameter, on sucrose acceptance, examining the three-way interactions with humidity and using the model formula:

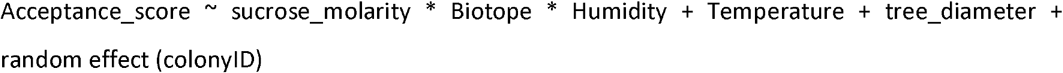

And examining the three-way interactions with temperature and using the model formula:

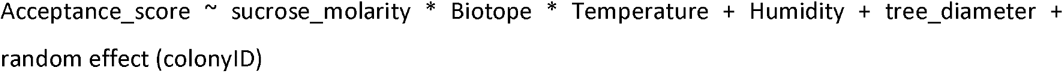

Finally, we explored the effect of tree characteristics on traffic. For this, we excluded data from three tree species which were only sampled once, leaving three tree species. We used the following GLM with a gaussian distribution:

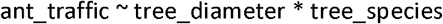

P values were later corrected for multiple testing using a Benjamini-Hochberg correction (Benjamini and Hochberg 1995). Interactions were analysed by subsetting: for instance, if the interaction food concentration * biome was significant, two subsets were made, each containing just one biome, and the analysis were rerun only including food concentration measure. For graphical visualization, raw data figures were created using ggplot2 (Wickham et al. 2020), while model predictions were visualized with ggeffect (Lüdecke 2018).

## Results

The complete raw data on which the analysis was based is provided in online supplement S1. The complete analysis document, with all code and analysis results, is provided in online supplement S2.

We found a significant interaction between sucrose concentration and habitat type on acceptance scores (X^2^ = 22.9 P < 0.0001). Overall, acceptance scores increased with sucrose concentration (X^2^ = 1649, P < 0.0001) and ants from urban habitats exhibited higher levels of sucrose acceptance compared to the rural counterparts (X^2^ = 16.27, P < 0.0001). Moreover, the increase in acceptance with rising sucrose concentration was more pronounced in urban populations than the rural ones. At the highest concentration tested (2M), acceptance level converged with no significant difference between urban and rural ant populations (see figure 1).

**Figure 1.**
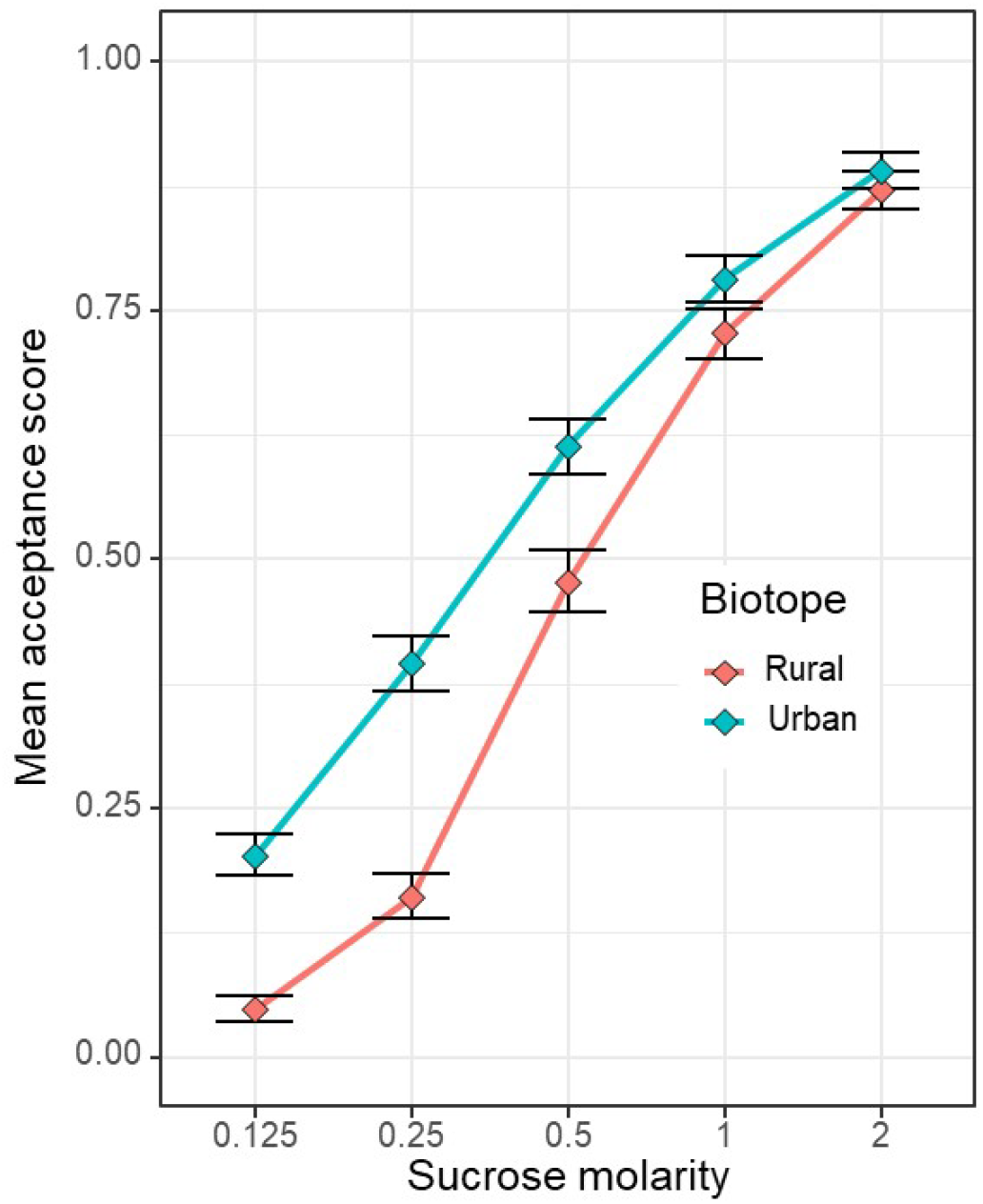
Mean acceptance scores of ants from rural or urban areas in response to a range of sucrose solution molarities Symbols are means, whiskers are bootstrapped 95% confidence intervals.

Humidity had no effect on sucrose acceptance, either alone or in a three-way interaction between humidity, sucrose concentration, and biotope (Humidity: X^2^ = 0.007, P = 0.93, three-way interaction: X^2^ = 0.068, P = 0.79). However, while temperature had no predictive power on sucrose acceptance alone (X^2^ = 1.51, P = 0.22), it did have influence on how ants in different biomes respond differentially to different sucrose molarities (three-way interaction, X^2^ = 8.38, P < 0.0038).

Exploring the three-way interaction by examining the predictions for the minimum, mean, and maximum temperatures (figure 2), we see that in urban environments at the lowest and highest temperatures, sucrose acceptance is reduced compared to at mean temperatures, especially for high sucrose levels.

**Figure 2.**
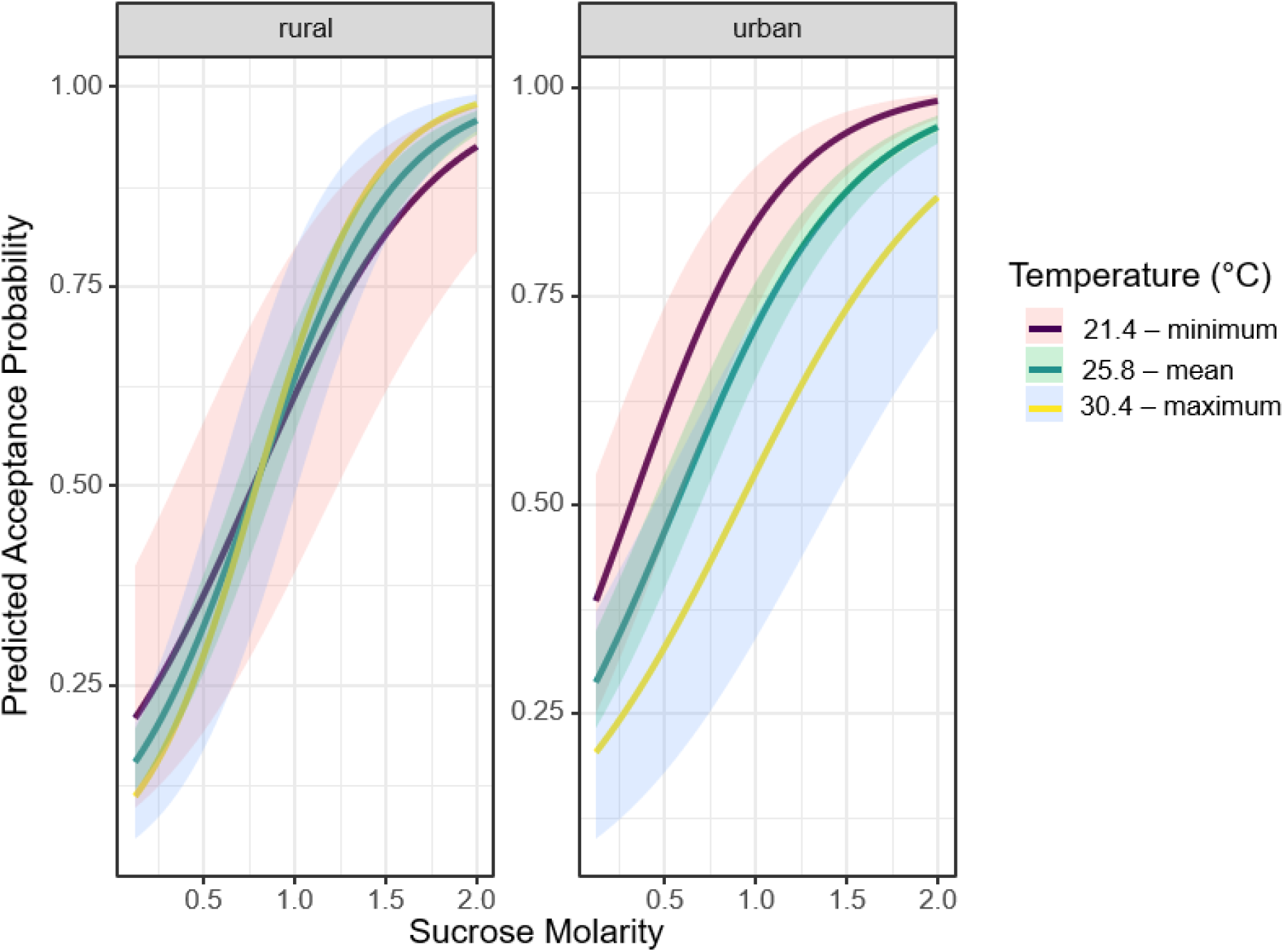
Predicted acceptance scores of ants from rural or urban areas in response to a range of sucrose solution molarities, at the minimum, maximum, and mean temperatures. Lines are glmm model predictions, ribbons are the 95% confidence interval of each prediction.

We found a positive correlation between tree diameter size and ant traffic (X^2^ = 8.6, P = 0.0033), significant differences in traffic between different tree species (X^2^ = 7.45, P = 0.024), and a non-significant interaction between species and diameter (X^2^ = 5.1, P = 0.078), in which increasing diameter strongly increases traffic on *Acer platanoides* but only slightly increases traffic *on Populus nigra* and *Tilia cordata* (see figure 3). Figure 3 shows the interaction – a plot showing the linear response without interaction can be found in the code and analysis supplement.

**Figure 3.**
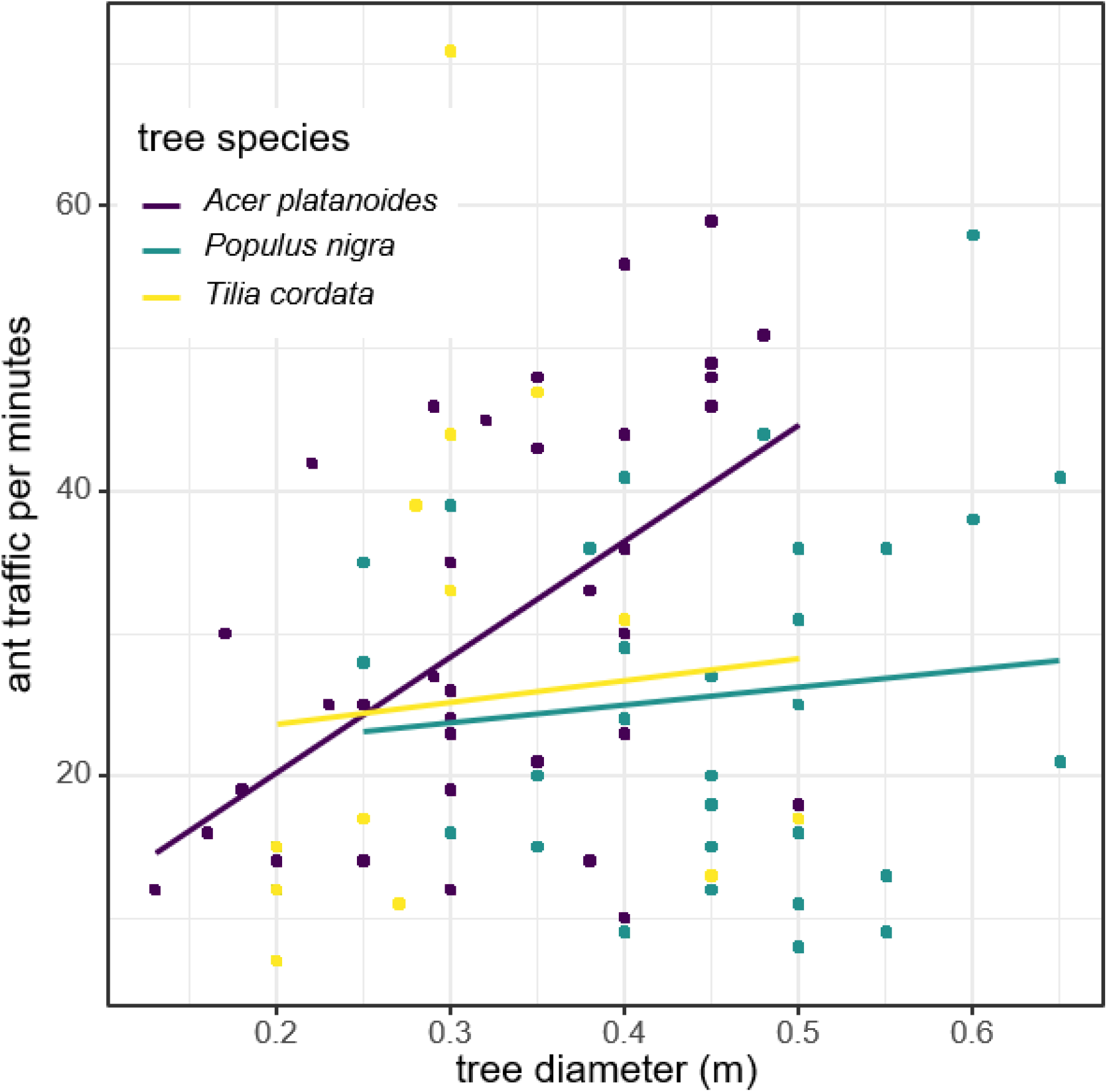
Relationship between tree diameter and ant traffic. Traffic increases on larger trees, especially in Norway Maple (*Acer palatanoides*). Lines are linear regressions, dots are individual values.

## Discussion

The sweeter a sucrose solution is, the more likely ants are to accept it, with ants generally rejecting low-molarity sucrose solutions (Wendt et al. 2019). However, we detected a marked difference in the response to low-molarity sucrose solutions between urban and rural populations: Urban ants exhibited a greater acceptance for these lower-quality resources.

Sugary substances are particularly attractive to many ant species, especially aphidicolous species such as *Lasius niger*, as they constitute their primary carbohydrate—and thus energy—source (Wilson and Eisner 1957; Markin 1970). Under natural conditions, ants primarily obtain carbohydrates from aphid honeydew, the composition of which varies significantly depending on factors such as soil chemistry, plant physiological state, and aphid species (Völkl et al. 1999).

One especially attractive component of honeydew for ants is melicitose (Cornelius et al. 1996; Detrain and Prieur 2014). The concentration of melicitose in honeydew is sensitive to environmental factors affecting the host plant’s physiological condition, often increasing under conditions of thermal or hydric stress (Seeburger et al. 2022). Given these factors, it is plausible that honeydew produced by sap-feeding insects in urban environments contains elevated levels of melicitose compared to that from rural areas. This is likely due to the fact that urban vegetation is often subject to greater thermal stress, driven by elevated ambient temperatures (Zhou et al., 2017), and reduced water availability, as impervious surfaces restrict soil infiltration and promote surface runoff. Then, ants - known for their sensitivity to subtle changes in the nutritional composition of food (Csata et al. 2020) – should be likely to detect and respond to changes in the concentration of carbohydrates in the honeydew. Indeed, Wendt and Czaczkes (2017) demonstrated that worker ants were willing to travel greater distances—thus expending more energy and increasing their risk of mortality—when the sugar concentration of a food source was higher. Furthermore, ants are capable of learning and forming expectations about food quality based on prior experience. For example, Wendt et al. (2019) found that ants accustomed to high-sugar diets were less inclined to accept food sources with lower sugar concentrations.

Based on all above information, it could be expected that urban ants will show reduced acceptance of low-sugar concentrations. Contrary to this expectation, but in line with previous published works, our study found that urban populations of *L. niger* were more willing to accept low concentrations of sugar than their rural counterparts, suggesting a lower quality of honeydew in urban environments. As an example of the nutritional constraints of urban areas, Garber et al. (2024) report that urban *L. niger* colonies defended aphid clusters more fiercely than their counterparts in rural areas, suggesting a potentially greater reliance on this resource due to increased food scarcity in cities. In line with these findings, Penick et al. (2015) also found that ants from urban areas fed on human waste, which is an abundant, energy-rich, and accessible dietary alternative in urban environments

In addition to general differences in resource acceptance, rural populations of *Lasius niger* exhibited a more pronounced increase in interest in sugary solutions as concentration levels rose. Previous studies have shown that urban plants are more susceptible to aphid infestations than their rural counterparts (Lubiarz et al. 2011; Mackos-Iwaszko et al. 2015), potentially providing ants with a greater abundance of honeydew. However, the nutritional quality of this honeydew may be compromised by the physiological stress experienced by urban plants. Our findings support this interpretation: urban ants showed a consistent interest in alternative sugar sources regardless of concentration, indicating a lower baseline quality of naturally available carbohydrates in urban environments. Consistent with the observations of Wendt et al. (2019), this pattern suggests that the more selective response of rural ants and their sharp increase in sugar consumption only at concentrations above 0.5 M might be linked to their access to higher-quality honeydew. Similarly, Trigos-Peral et al. (2024a) found that urban ant populations of *Lasius niger* in Warsaw, Poland, consumed more carbohydrate-rich (sugar-water) solutions and less protein than their rural counterparts. These habitat-related dietary shifts were interpreted as evidence of lower dietary quality in urban environments and were accompanied by physiological differences, with urban gynes exhibited lower body fat content.

Taken together, the results reported here and elsewhere support the potential of ants as bioindicators of dietary shifts and environmental quality. As previously noted, ants are widely recognized for their accuracy and reliability as indicators of environmental conditions, owing to their high sensitivity to habitat alterations and ecological disturbances (Andersen 1997). The findings of this study extend the known sensitivity of ants as a bioindicator group, demonstrating their potential to detect not only structural or abiotic changes in the environment (Trigos-Peral et al. 2024b) but also shifts in the nutritional landscape. This broadens their applicability across a wider range of ecological research. Furthermore, the methodology employed in this study is low-cost, easy to implement, rapid, and yields reliable results within a short timeframe, making it a practical tool for monitoring environmental quality, either by trained specialists or citizen scientists.

## Acknowledgements

TJC was supported by a Starter Grant from the European Research Council (CognitiveControl: 948181) and a Heisenberg grant from the Deutsche Forschungsgemeinschaft (DFG, German Research Foundation)—Projektnummer 462101190.

