## supplement S1 for "Urban *Lasius niger* ants more readily accept low concentration sucrose solution than rural ants": S1 - complete analysis code and output.html

Ants as Bioindicators - how level of urbanisation affects sucrose acceptance in Lasius niger ants


### Ants as Bioindicators - how level of urbanisation affects sucrose acceptance in Lasius niger ants

###### Analysis by Tomer J Czaczkes

#### 24.02.2025

In this study, ants in Urban and rural enviroments were offered
various sucrose concentrations as they walked towards established
feeding trees, and their acceptance of the sucrose was quantified.

### setting seed, activating packages

```
set.seed(123)
Sys.setlocale("LC_ALL", "en_US.UTF-8") # fixing degrees symbol problem

library(ggplot2)
library(ggeffects) # for marignal effect plots
library (emmeans)
```

```
## Welcome to emmeans.
## Caution: You lose important information if you filter this package's results.
## See '? untidy'
```

```
library(readxl)
library(dplyr)
```

```
## 
## Attaching package: 'dplyr'
```

```
## The following objects are masked from 'package:stats':
## 
##     filter, lag
```

```
## The following objects are masked from 'package:base':
## 
##     intersect, setdiff, setequal, union
```

```
library(glmmTMB)
library (car)
```

```
## Loading required package: carData
```

```
## 
## Attaching package: 'car'
```

```
## The following object is masked from 'package:dplyr':
## 
##     recode
```

```
library(stats) # for p.adjust, to do correction for multiple testing
library (eoffice)
```

### importing data, set working directory

```
data <- read_excel("data_2024_unblinded.xlsx")
only_common_trees <- subset(data,
                        !(tree_species %in% c("Acer campestre", "Juglans regia", "Quercus robur", "Ulmus")))
#first only the first ant
onlyfirstant_onlycommontrees <- subset(only_common_trees, number_of_observation == 1)

#now only one of the molarities, because only after each full round was the traffic rechecked. Randomly choosing 0.5
much_reduced <- subset(onlyfirstant_onlycommontrees, sucrose_molarity == 0.5)
```

quick check of the data, see if everything is alright

```
str (data)
```

```
## tibble [9,000 × 15] (S3: tbl_df/tbl/data.frame)
##  $ Data                       : POSIXct[1:9000], format: "2024-08-15" "2024-08-15" ...
##  $ Coordinates                : chr [1:9000] "50.56674545655925, 30.21840408441833" "50.56674545655925, 30.21840408441833" "50.56674545655925, 30.21840408441833" "50.56674545655925, 30.21840408441833" ...
##  $ Location                   : chr [1:9000] "Bucha" "Bucha" "Bucha" "Bucha" ...
##  $ Temperature                : num [1:9000] 26.8 26.8 26.8 26.8 26.8 26.8 26.8 26.8 26.8 26.8 ...
##  $ Humidity                   : num [1:9000] 47 47 47 47 47 47 47 47 47 47 ...
##  $ colonyID                   : chr [1:9000] "C46" "C46" "C46" "C46" ...
##  $ ant_traffic                : num [1:9000] 31 31 31 31 31 31 31 31 31 31 ...
##  $ Biotope                    : chr [1:9000] "rural" "rural" "rural" "rural" ...
##  $ sucrose_molarity           : num [1:9000] 0.125 0.125 0.125 0.125 0.125 0.125 0.125 0.125 0.125 0.125 ...
##  $ number_of_observation      : num [1:9000] 1 2 3 4 5 6 7 8 9 10 ...
##  $ acceptance_score           : num [1:9000] 1 1 1 1 0.5 0 0 0 0 0.5 ...
##  $ acceptance_score_halfisone : num [1:9000] 1 1 1 1 1 0 0 0 0 1 ...
##  $ acceptance_score_halfiszero: num [1:9000] 1 1 1 1 0 0 0 0 0 0 ...
##  $ tree_species               : chr [1:9000] "Populus nigra" "Populus nigra" "Populus nigra" "Populus nigra" ...
##  $ tree_diameter              : num [1:9000] 0.5 0.5 0.5 0.5 0.5 0.5 0.5 0.5 0.5 0.5 ...
```

```
summary (data)
```

```
##       Data                     Coordinates          Location        
##  Min.   :2024-08-14 00:00:00   Length:9000        Length:9000       
##  1st Qu.:2024-08-17 00:00:00   Class :character   Class :character  
##  Median :2024-08-20 12:00:00   Mode  :character   Mode  :character  
##  Mean   :2024-08-20 12:00:00                                        
##  3rd Qu.:2024-08-24 00:00:00                                        
##  Max.   :2024-08-27 00:00:00                                        
##                                                                     
##   Temperature       Humidity       colonyID          ant_traffic   
##  Min.   :21.40   Min.   :29.00   Length:9000        Min.   : 7.00  
##  1st Qu.:24.40   1st Qu.:40.00   Class :character   1st Qu.:15.00  
##  Median :25.55   Median :45.00   Mode  :character   Median :24.50  
##  Mean   :25.85   Mean   :46.14                      Mean   :27.28  
##  3rd Qu.:27.40   3rd Qu.:49.00                      3rd Qu.:38.00  
##  Max.   :30.40   Max.   :84.00                      Max.   :71.00  
##                                                                    
##    Biotope          sucrose_molarity number_of_observation acceptance_score
##  Length:9000        Min.   :0.125    Min.   : 1.00         Min.   :0.000   
##  Class :character   1st Qu.:0.250    1st Qu.: 5.75         1st Qu.:0.000   
##  Mode  :character   Median :0.500    Median :10.50         Median :0.500   
##                     Mean   :0.775    Mean   :10.50         Mean   :0.517   
##                     3rd Qu.:1.000    3rd Qu.:15.25         3rd Qu.:1.000   
##                     Max.   :2.000    Max.   :20.00         Max.   :1.000   
##                                                            NA's   :1       
##  acceptance_score_halfisone acceptance_score_halfiszero tree_species      
##  Min.   :0.0000             Min.   :0.0000              Length:9000       
##  1st Qu.:0.0000             1st Qu.:0.0000              Class :character  
##  Median :1.0000             Median :0.0000              Mode  :character  
##  Mean   :0.5952             Mean   :0.4388                                
##  3rd Qu.:1.0000             3rd Qu.:1.0000                                
##  Max.   :1.0000             Max.   :1.0000                                
##  NA's   :1                  NA's   :1                                     
##  tree_diameter   
##  Min.   :0.1300  
##  1st Qu.:0.2900  
##  Median :0.3500  
##  Mean   :0.3632  
##  3rd Qu.:0.4500  
##  Max.   :0.6500  
##
```

### Figures

#### Effect of sucrose concentration on acceptance

We have 0.5s in the data (half accept). It will be much easier to do
stats on the data if it is binomial. Let’s see how different the
stringent and lenient data coding look:

```
## `summarise()` has grouped output by 'Biotope'. You can override using the
## `.groups` argument.
## `summarise()` has grouped output by 'Biotope'. You can override using the
## `.groups` argument.
## `summarise()` has grouped output by 'Biotope'. You can override using the
## `.groups` argument.
```

It doesn’t change the story at all: looks like the 0.5s are pretty well
distributed. turing 0.5 into 1 emphasies differences at lower molarities
and makes differences smaller at higher molarities. Hard to tell which
is best to use, so let’s just stick with the raw data (0.5 as 0.5) for
now, as a binomial distribution family should still be ok.

#### effect of tempreture, humidity, tree diameter, and traffic on trunk trail?

```
#first temp
ggplot(data, aes(x = Temperature, y = acceptance_score, fill = Biotope)) +
geom_point() + 
geom_smooth (n = 5) +
  ylab("Mean acceptance score") +
  xlab("Temperature") +
  theme_bw(15)
```

```
## `geom_smooth()` using method = 'gam' and formula = 'y ~ s(x, bs = "cs")'
```

```
#now humidity
ggplot(data, aes(x = Humidity, y = acceptance_score, fill = Biotope)) +
geom_point() + 
geom_smooth (n = 5) +
  ylab("Mean acceptance score") +
  xlab("Humidity") +
  theme_bw(15)
```

```
## `geom_smooth()` using method = 'gam' and formula = 'y ~ s(x, bs = "cs")'
```

```
#now tree diameter 
ggplot(data, aes(x = tree_diameter, y = acceptance_score, fill = Biotope)) +
geom_point() + 
geom_smooth (n = 5) +
  ylab("Mean acceptance score") +
  xlab("tree diameter") +
  theme_bw(15)
```

```
## `geom_smooth()` using method = 'gam' and formula = 'y ~ s(x, bs = "cs")'
```

```
#finally ant traffic
ggplot(data, aes(x = ant_traffic, y = acceptance_score, fill = Biotope)) +
geom_point() + 
geom_smooth (n = 5) +
  ylab("Mean acceptance score") +
  xlab("ant traffic per minutes") +
  theme_bw(15)
```

```
## `geom_smooth()` using method = 'gam' and formula = 'y ~ s(x, bs = "cs")'
```

No clear effects of temp, humidity, or tree diameter, but ant traffic
has an odd effect, wherein in urban enviroments acceptance goes up at
higher traffic levels. Are traffic levels higher in Urban enviroments
too?

```
#finally ant traffic
ggplot(data, aes(x = Biotope, y = ant_traffic, fill = Biotope)) +
  geom_violin(alpha=0.5)+ 
stat_summary(fun.data = "mean_cl_boot", geom = "errorbar", width = 0.5) +
stat_summary(fun.y = "mean", geom = "point", shape = 23, size = 3) +
  ylab("Ant traffic") +
  xlab("Biotope") +
  theme_bw(15)
```

```
ggplot(data, aes(x = ant_traffic, fill = Biotope)) +
  geom_histogram(position = "identity", alpha = 0.5, bins = 20) +
  scale_fill_manual(values = c("red","blue")) +  # Adjust colors as needed
  labs(title = "Distribution of Ant Traffic by Biotope",
       x = "Ant Traffic",
       y = "Frequency") +
  theme_minimal()
```

No, it looks actually like ant traffic is lower in Urban enviroments
overall. No obvious differences in distribution - just a more higher
traffic paths in rural enviroments. No real way of telling why - true
effect? Sampling bias? Impossible to say.

#### Tree characteristics

##### tree characteristics and ant traffic

How does tree diameter and species affect traffic

```
# effect of tree diameter on ant traffic
ggplot(data, aes(x = tree_diameter, y = ant_traffic)) +
geom_point() + 
geom_smooth (n = 2) +
  ylab("ant traffic per minutes") +
  xlab("tree diameter (m)") +
  theme_bw(15)
```

```
## `geom_smooth()` using method = 'gam' and formula = 'y ~ s(x, bs = "cs")'
```

```
# effect of tree species on ant traffic 
ggplot(data, aes(x = tree_species, y = ant_traffic)) +
  geom_violin(alpha=0.5)+ 
  geom_jitter(alpha = 0.1) +
stat_summary(fun.data = "mean_cl_boot", geom = "errorbar", width = 0.5) +
stat_summary(fun.y = "mean", geom = "point", shape = 23, size = 3, fill = "red") +
  ylab("Ant traffic") +
  xlab("Tree species") +
  theme_bw(15) + 
  theme(axis.text.x = element_text(angle = 45, hjust = 1))
```

It seems that, nicely but unsurprisingly, we have higher traffic on
larger trees. We will analyse this properly below. Tree species is
harder to work with, since 3 species are only represented by one
tree.

Let’s remove the three underrepresented trees, and then look
again:

##### interaction of tree species and tree diameter with ant traffic

```
# first remove the three uncommon species
only_common_trees <- subset(data,
                        !(tree_species %in% c("Acer campestre", "Juglans regia", "Quercus robur", "Ulmus")))

# now an interaction plot
diameter_plot <- ggplot(much_reduced, aes(x = tree_diameter, y = ant_traffic, color = tree_species)) +
  geom_point(alpha = 0.6, show.legend = FALSE) + # Individual value points, adjust alpha for transparency
  geom_smooth(method = "lm", se = FALSE) + # Regression lines, se=TRUE adds confidence interval
  ylab("ant traffic per minutes") +
  xlab("tree diameter (m)") +
  theme_bw(15) + 
   scale_color_viridis_d(option = "D", name = "tree species") +# makes the plot ok for people with color blindness
 theme(legend.text = element_text(face = "italic"))

print (diameter_plot)
```

```
## `geom_smooth()` using formula = 'y ~ x'
```

ok, it looks like there is a strong correlation between tree size and
traffic for Acer platanoides, but not for Poplars and Tilia.

#### exploring colonyID

Finally, I would like to see how much variation we really have
between colonies.

```
ggplot(data, aes(x = as.factor(colonyID), y = acceptance_score, fill = Biotope)) +
stat_summary(fun.data = "mean_cl_boot", geom = "errorbar", width = 0.5) +
stat_summary(fun.y = "mean", geom = "point", shape = 23, size = 3) +
  ylab("acceptance") +
  xlab("colony") +
  theme_bw(15)
```

```
## Warning: Removed 1 row containing non-finite outside the scale range (`stat_summary()`).
## Removed 1 row containing non-finite outside the scale range (`stat_summary()`).
```

Ok, yes, I believe that Urban enviroments really do have higher
acceptances overall.

### statistical analysis

note - analysis of temperature, traffic, tree diameter is relegated
to the bottom of the document, as these are uninteresting, were excluded
in exploration, and are included only for completeness.

#### top level analysis

```
m1 = glmmTMB (acceptance_score ~ sucrose_molarity * Biotope
            +  (1|colonyID),  #random effects - ant ID nested within colony ID
            family = binomial,   #distribution family
            data=data    #what data to use
)


anova_results <- Anova (m1)
print (anova_results)
```

```
## Analysis of Deviance Table (Type II Wald chisquare tests)
## 
## Response: acceptance_score
##                             Chisq Df Pr(>Chisq)    
## sucrose_molarity         1649.116  1  < 2.2e-16 ***
## Biotope                    16.269  1  5.495e-05 ***
## sucrose_molarity:Biotope   22.933  1  1.677e-06 ***
## ---
## Signif. codes:  0 '***' 0.001 '**' 0.01 '*' 0.05 '.' 0.1 ' ' 1
```

```
# Extract the p-values from the Anova results
p_values <- anova_results$"Pr(>Chisq)"

# Perform Benjamini-Hochberg correction
adjusted_p_values <- p.adjust(p_values, method = "BH")

# Combine the results into a data frame
adjusted_results <- cbind(anova_results, Adjusted_P_Value = adjusted_p_values)

# Print the adjusted results
print(adjusted_results)
```

```
##                               Chisq Df   Pr(>Chisq) Adjusted_P_Value
## sucrose_molarity         1649.11558  1 0.000000e+00     0.000000e+00
## Biotope                    16.26903  1 5.495489e-05     5.495489e-05
## sucrose_molarity:Biotope   22.93335  1 1.677169e-06     2.515753e-06
```

Ok, nice clear result: interaction between food concentration and
Biotope, and also food concentration and biotope are themselves highly
significant.

##### sugar \* Biome effect: understanding interactions

let’s try to understand the interaction by looking at marginal
effects

```
ggeffect_sugarplot <- plot(ggeffect(m1, c("sucrose_molarity", "Biotope")))
```

```
## You are calculating adjusted predictions on the population-level (i.e.
##   `type = "fixed"`) for a *generalized* linear mixed model.
##   This may produce biased estimates due to Jensen's inequality. Consider
##   setting `bias_correction = TRUE` to correct for this bias.
##   See also the documentation of the `bias_correction` argument.
```

```
ggeffect_sugarplot <- ggeffect_sugarplot +
  theme_bw(15) +
  coord_cartesian(ylim = c(0,1)) + 
  ylab("Predicted acceptance score") +
  xlab("Sucrose molarity") +
  labs(title = NULL)

plot (ggeffect_sugarplot)
```

Ok, so it looks like sugar concentration strongly correlates with
acceptance in both enviroments, but the increasing sucrose concentration
has less of an effect at higher concentrations in urban enviroments. At
highest sucrose concentration (2m) acceptances scores are the same

```
Urban_sugar_data <- subset (data, Biotope == "urban")
rural_sugar_data <- subset (data, Biotope == "rural")

Urban_model_sugar = glmmTMB (acceptance_score ~ sucrose_molarity
            +  (1|colonyID),  #random effects - ant ID nested within colony ID
            family = binomial,   #distribution family
            data=Urban_sugar_data    #what data to use
)

rural_model_sugar = glmmTMB (acceptance_score ~ sucrose_molarity
            +  (1|colonyID),  #random effects - ant ID nested within colony ID
            family = binomial,   #distribution family
            data=rural_sugar_data    #what data to use
)

## urban model
anova_results <- Anova (Urban_model_sugar)
print (anova_results)
```

```
## Analysis of Deviance Table (Type II Wald chisquare tests)
## 
## Response: acceptance_score
##                   Chisq Df Pr(>Chisq)    
## sucrose_molarity 702.87  1  < 2.2e-16 ***
## ---
## Signif. codes:  0 '***' 0.001 '**' 0.01 '*' 0.05 '.' 0.1 ' ' 1
```

```
# Extract the p-values from the Anova results
p_values <- anova_results$"Pr(>Chisq)"

# Perform Benjamini-Hochberg correction
adjusted_p_values <- p.adjust(p_values, method = "BH")

# Combine the results into a data frame
adjusted_results <- cbind(anova_results, Adjusted_P_Value = adjusted_p_values)

# Print the adjusted results
print(adjusted_results)
```

```
##                     Chisq Df    Pr(>Chisq) Adjusted_P_Value
## sucrose_molarity 702.8665  1 7.118055e-155    7.118055e-155
```

```
summary (Urban_model_sugar)
```

```
##  Family: binomial  ( logit )
## Formula:          acceptance_score ~ sucrose_molarity + (1 | colonyID)
## Data: Urban_sugar_data
## 
##       AIC       BIC    logLik -2*log(L)  df.resid 
##    4707.6    4726.8   -2350.8    4701.6      4496 
## 
## Random effects:
## 
## Conditional model:
##  Groups   Name        Variance Std.Dev.
##  colonyID (Intercept) 0.663    0.8143  
## Number of obs: 4499, groups:  colonyID, 45
## 
## Conditional model:
##                  Estimate Std. Error z value Pr(>|z|)    
## (Intercept)      -1.06886    0.13403  -7.975 1.53e-15 ***
## sucrose_molarity  2.10957    0.07957  26.512  < 2e-16 ***
## ---
## Signif. codes:  0 '***' 0.001 '**' 0.01 '*' 0.05 '.' 0.1 ' ' 1
```

```
## rural model
anova_results <- Anova (rural_model_sugar)
print (anova_results)
```

```
## Analysis of Deviance Table (Type II Wald chisquare tests)
## 
## Response: acceptance_score
##                   Chisq Df Pr(>Chisq)    
## sucrose_molarity 966.63  1  < 2.2e-16 ***
## ---
## Signif. codes:  0 '***' 0.001 '**' 0.01 '*' 0.05 '.' 0.1 ' ' 1
```

```
# Extract the p-values from the Anova results
p_values <- anova_results$"Pr(>Chisq)"

# Perform Benjamini-Hochberg correction
adjusted_p_values <- p.adjust(p_values, method = "BH")

# Combine the results into a data frame
adjusted_results <- cbind(anova_results, Adjusted_P_Value = adjusted_p_values)

# Print the adjusted results
print(adjusted_results)
```

```
##                     Chisq Df    Pr(>Chisq) Adjusted_P_Value
## sucrose_molarity 966.6309  1 3.218251e-212    3.218251e-212
```

```
summary (rural_model_sugar)
```

```
##  Family: binomial  ( logit )
## Formula:          acceptance_score ~ sucrose_molarity + (1 | colonyID)
## Data: rural_sugar_data
## 
##       AIC       BIC    logLik -2*log(L)  df.resid 
##    4204.6    4223.9   -2099.3    4198.6      4497 
## 
## Random effects:
## 
## Conditional model:
##  Groups   Name        Variance Std.Dev.
##  colonyID (Intercept) 0.668    0.8173  
## Number of obs: 4500, groups:  colonyID, 45
## 
## Conditional model:
##                  Estimate Std. Error z value Pr(>|z|)    
## (Intercept)      -2.12675    0.13961  -15.23   <2e-16 ***
## sucrose_molarity  2.66193    0.08562   31.09   <2e-16 ***
## ---
## Signif. codes:  0 '***' 0.001 '**' 0.01 '*' 0.05 '.' 0.1 ' ' 1
```

Much as the figures suggest: intercept for rural is lower, but slope
is higher

#### Analysis of temperature, humidity, tree diameter:

```
m_TempHumidTree = glmmTMB (acceptance_score ~ sucrose_molarity * Biotope + Humidity + Temperature + tree_diameter
            +  (1|colonyID),  #random effects - ant ID nested within colony ID
            family = binomial,   #distribution family
            data=data    #what data to use
)


anova_results <- Anova (m_TempHumidTree)
print (anova_results)
```

```
## Analysis of Deviance Table (Type II Wald chisquare tests)
## 
## Response: acceptance_score
##                              Chisq Df Pr(>Chisq)    
## sucrose_molarity         1649.2167  1  < 2.2e-16 ***
## Biotope                     6.0887  1    0.01361 *  
## Humidity                    0.0066  1    0.93508    
## Temperature                 2.3272  1    0.12713    
## tree_diameter               0.9175  1    0.33813    
## sucrose_molarity:Biotope   22.5739  1  2.022e-06 ***
## ---
## Signif. codes:  0 '***' 0.001 '**' 0.01 '*' 0.05 '.' 0.1 ' ' 1
```

```
# Extract the p-values from the Anova results
p_values <- anova_results$"Pr(>Chisq)"

# Perform Benjamini-Hochberg correction
adjusted_p_values <- p.adjust(p_values, method = "BH")

# Combine the results into a data frame
adjusted_results <- cbind(anova_results, Adjusted_P_Value = adjusted_p_values)

# Print the adjusted results
print(adjusted_results)
```

```
##                                 Chisq Df   Pr(>Chisq) Adjusted_P_Value
## sucrose_molarity         1.649217e+03  1 0.000000e+00     0.000000e+00
## Biotope                  6.088665e+00  1 1.360519e-02     2.721037e-02
## Humidity                 6.634264e-03  1 9.350833e-01     9.350833e-01
## Temperature              2.327249e+00  1 1.271264e-01     1.906895e-01
## tree_diameter            9.175041e-01  1 3.381312e-01     4.057574e-01
## sucrose_molarity:Biotope 2.257389e+01  1 2.022143e-06     6.066428e-06
```

##### three way interactions: Analysis of temperature, humidity, tree diameter:

###### sucrose*biotope*humidity

```
m_threeway_humid = glmmTMB (acceptance_score ~ sucrose_molarity * Biotope * Humidity + Temperature + tree_diameter
            +  (1|colonyID),  #random effects - ant ID nested within colony ID
            family = binomial,   #distribution family
            data=data    #what data to use
)


anova_results <- Anova (m_threeway_humid)
print (anova_results)
```

```
## Analysis of Deviance Table (Type II Wald chisquare tests)
## 
## Response: acceptance_score
##                                       Chisq Df Pr(>Chisq)    
## sucrose_molarity                  1646.9767  1  < 2.2e-16 ***
## Biotope                              6.3039  1    0.01205 *  
## Humidity                             0.0075  1    0.93100    
## Temperature                          1.5155  1    0.21829    
## tree_diameter                        1.9038  1    0.16765    
## sucrose_molarity:Biotope            23.9661  1  9.804e-07 ***
## sucrose_molarity:Humidity            0.9532  1    0.32890    
## Biotope:Humidity                     2.2575  1    0.13297    
## sucrose_molarity:Biotope:Humidity    0.0680  1    0.79432    
## ---
## Signif. codes:  0 '***' 0.001 '**' 0.01 '*' 0.05 '.' 0.1 ' ' 1
```

```
# Extract the p-values from the Anova results
p_values <- anova_results$"Pr(>Chisq)"

# Perform Benjamini-Hochberg correction
adjusted_p_values <- p.adjust(p_values, method = "BH")

# Combine the results into a data frame
adjusted_results <- cbind(anova_results, Adjusted_P_Value = adjusted_p_values)

# Print the adjusted results
print(adjusted_results)
```

```
##                                          Chisq Df   Pr(>Chisq) Adjusted_P_Value
## sucrose_molarity                  1.646977e+03  1 0.000000e+00     0.000000e+00
## Biotope                           6.303950e+00  1 1.204693e-02     3.614078e-02
## Humidity                          7.498060e-03  1 9.309964e-01     9.309964e-01
## Temperature                       1.515546e+00  1 2.182947e-01     3.274421e-01
## tree_diameter                     1.903839e+00  1 1.676492e-01     3.017686e-01
## sucrose_molarity:Biotope          2.396615e+01  1 9.804458e-07     4.412006e-06
## sucrose_molarity:Humidity         9.532282e-01  1 3.288989e-01     4.228701e-01
## Biotope:Humidity                  2.257504e+00  1 1.329682e-01     2.991785e-01
## sucrose_molarity:Biotope:Humidity 6.796698e-02  1 7.943202e-01     8.936102e-01
```

###### sucrose*biotope*temperature

```
m_threeway_temp = glmmTMB (acceptance_score ~ sucrose_molarity * Biotope * Temperature + Humidity + tree_diameter
            +  (1|colonyID),  #random effects - ant ID nested within colony ID
            family = binomial,   #distribution family
            data=data    #what data to use
)


anova_results <- Anova (m_threeway_temp)
print (anova_results)
```

```
## Analysis of Deviance Table (Type II Wald chisquare tests)
## 
## Response: acceptance_score
##                                          Chisq Df Pr(>Chisq)    
## sucrose_molarity                     1636.7038  1  < 2.2e-16 ***
## Biotope                                 5.9592  1   0.014640 *  
## Temperature                             2.3450  1   0.125690    
## Humidity                                0.0036  1   0.952307    
## tree_diameter                           0.9157  1   0.338597    
## sucrose_molarity:Biotope               20.8565  1   4.95e-06 ***
## sucrose_molarity:Temperature            0.0234  1   0.878420    
## Biotope:Temperature                     1.0843  1   0.297741    
## sucrose_molarity:Biotope:Temperature    8.3808  1   0.003792 ** 
## ---
## Signif. codes:  0 '***' 0.001 '**' 0.01 '*' 0.05 '.' 0.1 ' ' 1
```

```
# Extract the p-values from the Anova results
p_values <- anova_results$"Pr(>Chisq)"

# Perform Benjamini-Hochberg correction
adjusted_p_values <- p.adjust(p_values, method = "BH")

# Combine the results into a data frame
adjusted_results <- cbind(anova_results, Adjusted_P_Value = adjusted_p_values)

# Print the adjusted results
print(adjusted_results)
```

```
##                                             Chisq Df   Pr(>Chisq)
## sucrose_molarity                     1.636704e+03  1 0.000000e+00
## Biotope                              5.959241e+00  1 1.464034e-02
## Temperature                          2.344950e+00  1 1.256896e-01
## Humidity                             3.577285e-03  1 9.523066e-01
## tree_diameter                        9.157352e-01  1 3.385973e-01
## sucrose_molarity:Biotope             2.085651e+01  1 4.950075e-06
## sucrose_molarity:Temperature         2.340059e-02  1 8.784200e-01
## Biotope:Temperature                  1.084286e+00  1 2.977407e-01
## sucrose_molarity:Biotope:Temperature 8.380794e+00  1 3.792068e-03
##                                      Adjusted_P_Value
## sucrose_molarity                         0.000000e+00
## Biotope                                  3.294076e-02
## Temperature                              2.262412e-01
## Humidity                                 9.523066e-01
## tree_diameter                            4.353393e-01
## sucrose_molarity:Biotope                 2.227534e-05
## sucrose_molarity:Temperature             9.523066e-01
## Biotope:Temperature                      4.353393e-01
## sucrose_molarity:Biotope:Temperature     1.137620e-02
```

We find a significant 3 way interaction. Let’s understand it using
emmeans:

It’s complex and long code - let’s let Gemini do the coding for
me!

```
# Assuming 'data' is the original data frame used to fit m_threeway_temp

# For the X-axis (sucrose_molarity): create a sequence of values
sucrose_molarity_seq <- seq(min(data$sucrose_molarity, na.rm = TRUE),
                            max(data$sucrose_molarity, na.rm = TRUE),
                            length.out = 50) # 50 points for a smooth line

# For the 'lines' (Temperature): choose 3 representative levels (e.g., min, mean, max)
# Using 'na.rm=TRUE' in case of NAs in your original data
temp_levels <- c(min(data$Temperature, na.rm = TRUE),
                 mean(data$Temperature, na.rm = TRUE),
                 max(data$Temperature, na.rm = TRUE))

# Alternatively, for robustness against outliers, use quantiles:
# temp_levels <- quantile(data$Temperature, probs = c(0.1, 0.5, 0.9), na.rm = TRUE)

# Convert to a data frame for easier handling if needed (though emmeans handles lists)
temp_levels_df <- data.frame(Temperature = temp_levels)

# For Biotope, emmeans will automatically use its two levels.

# Generate predictions
emm_predictions <- emmeans(m_threeway_temp,
                           specs = ~ sucrose_molarity * Temperature * Biotope,
                           at = list(sucrose_molarity = sucrose_molarity_seq,
                                     Temperature = temp_levels),
                           type = "response") # IMPORTANT: 'type = "response"' for probabilities

# Convert the emmeans object to a data frame for ggplot2
emm_df <- as.data.frame(emm_predictions)

emm_df$Temperature_f <- factor(round(emm_df$Temperature, 1)) # Round to 1 decimal place


temp_biome_sucrose_interactionplot <- ggplot(emm_df, aes(x = sucrose_molarity, y = prob, # 'prob' is the predicted probability
                   color = Temperature_f, fill = Temperature_f)) +
  geom_line(linewidth = 1.2) + # Draw the predicted lines
  geom_ribbon(aes(ymin = asymp.LCL, ymax = asymp.UCL), alpha = 0.2, linetype = 0) + # Add confidence intervals
  facet_wrap(~ Biotope, scales = "free_y") + # Separate panels for each Biotope level
                                            # scales="free_y" if probability ranges differ
  labs(
    x = "Sucrose Molarity",
    y = "Predicted Acceptance Probability",
    color = "Temperature (°C)" # Legend title for temperature levels
  ) +
  theme_bw(15) + # Your desired theme
  theme(legend.position = "right") +
  scale_color_viridis_d(option = "D") # Colorblind-friendly palette

print (temp_biome_sucrose_interactionplot)
```

interaction mostly due to in urban environments sucrose acceptance is
lower in more extreme temperatures (high and low), which temperature
doesn’t really affect acceptance in rural environments.

#### traffic analysis

```
# NOTE we will remove the three trees only sampled once for this analysis.
# NOTE cannot have colony ID as a random effect, since it's not well distributed around the trees. putting in Biome. Not ideal, but it lets me run the analysis

# note we need to subset the data because we have 20 repeats of each measure.
#and  only one of the molarities, because only after each full round was the traffic rechecked. Randomly choosing 0.5
# hence using "much reduced" dataset


traffic_treediameter  <- glm(ant_traffic ~ tree_diameter * tree_species,
                 family = gaussian, 
                 data = much_reduced)


anova_results <- Anova (traffic_treediameter)
print (anova_results)
```

```
## Analysis of Deviance Table (Type II tests)
## 
## Response: ant_traffic
##                            LR Chisq Df Pr(>Chisq)   
## tree_diameter                8.6073  1   0.003348 **
## tree_species                 7.4473  2   0.024145 * 
## tree_diameter:tree_species   5.0999  2   0.078087 . 
## ---
## Signif. codes:  0 '***' 0.001 '**' 0.01 '*' 0.05 '.' 0.1 ' ' 1
```

```
# Extract the p-values from the Anova results
p_values <- anova_results$"Pr(>Chisq)"

# Perform Benjamini-Hochberg correction
adjusted_p_values <- p.adjust(p_values, method = "BH")

# Combine the results into a data frame
adjusted_results <- cbind(anova_results, Adjusted_P_Value = adjusted_p_values)

# Print the adjusted results
print(adjusted_results)
```

```
##                            LR Chisq Df  Pr(>Chisq) Adjusted_P_Value
## tree_diameter              8.607336  1 0.003348116       0.01004435
## tree_species               7.447322  2 0.024145408       0.03621811
## tree_diameter:tree_species 5.099861  2 0.078087073       0.07808707
```

```
# overall summary with effects and interactions
summary (traffic_treediameter)
```

```
## 
## Call:
## glm(formula = ant_traffic ~ tree_diameter * tree_species, family = gaussian, 
##     data = much_reduced)
## 
## Coefficients:
##                                         Estimate Std. Error t value Pr(>|t|)
## (Intercept)                                3.983      7.546   0.528 0.599060
## tree_diameter                             81.287     22.314   3.643 0.000477
## tree_speciesPopulus nigra                 16.006     13.370   1.197 0.234757
## tree_speciesTilia cordata                 16.560     14.632   1.132 0.261128
## tree_diameter:tree_speciesPopulus nigra  -68.746     32.493  -2.116 0.037482
## tree_diameter:tree_speciesTilia cordata  -65.902     45.080  -1.462 0.147692
##                                            
## (Intercept)                                
## tree_diameter                           ***
## tree_speciesPopulus nigra                  
## tree_speciesTilia cordata                  
## tree_diameter:tree_speciesPopulus nigra *  
## tree_diameter:tree_speciesTilia cordata    
## ---
## Signif. codes:  0 '***' 0.001 '**' 0.01 '*' 0.05 '.' 0.1 ' ' 1
## 
## (Dispersion parameter for gaussian family taken to be 184.0855)
## 
##     Null deviance: 17734  on 85  degrees of freedom
## Residual deviance: 14727  on 80  degrees of freedom
## AIC: 700.36
## 
## Number of Fisher Scoring iterations: 2
```

### Session information - R version, packages, etc

Many thanks to all package creators!

```
## R version 4.5.0 (2025-04-11 ucrt)
## Platform: x86_64-w64-mingw32/x64
## Running under: Windows 10 x64 (build 19045)
## 
## Matrix products: default
##   LAPACK version 3.12.1
## 
## locale:
## [1] LC_COLLATE=en_US.UTF-8  LC_CTYPE=en_US.UTF-8    LC_MONETARY=en_US.UTF-8
## [4] LC_NUMERIC=C            LC_TIME=en_US.UTF-8    
## 
## time zone: Europe/Berlin
## tzcode source: internal
## 
## attached base packages:
## [1] stats     graphics  grDevices utils     datasets  methods   base     
## 
## other attached packages:
## [1] eoffice_0.2.2   car_3.1-3       carData_3.0-5   glmmTMB_1.1.11 
## [5] dplyr_1.1.4     readxl_1.4.5    emmeans_1.11.1  ggeffects_2.3.0
## [9] ggplot2_3.5.2  
## 
## loaded via a namespace (and not attached):
##   [1] DBI_1.2.3               Rdpack_2.6.4            gridExtra_2.3          
##   [4] sandwich_3.1-1          rlang_1.1.6             magrittr_2.0.3         
##   [7] multcomp_1.4-28         compiler_4.5.0          mgcv_1.9-1             
##  [10] systemfonts_1.2.3       vctrs_0.6.5             devEMF_4.5-1           
##  [13] stringr_1.5.1           pkgconfig_2.0.3         fastmap_1.2.0          
##  [16] backports_1.5.0         magick_2.8.6            labeling_0.4.3         
##  [19] rmarkdown_2.29          nloptr_2.2.1            ragg_1.4.0             
##  [22] purrr_1.0.4             xfun_0.52               cachem_1.1.0           
##  [25] jsonlite_2.0.0          uuid_1.2-1              cluster_2.1.8.1        
##  [28] broom_1.0.8             R6_2.6.1                R.devices_2.17.2       
##  [31] stringi_1.8.7           bslib_0.9.0             RColorBrewer_1.1-3     
##  [34] rpart_4.1.24            boot_1.3-31             jquerylib_0.1.4        
##  [37] cellranger_1.1.0        numDeriv_2016.8-1.1     estimability_1.5.1     
##  [40] Rcpp_1.0.14             knitr_1.50              zoo_1.8-14             
##  [43] base64enc_0.1-3         R.utils_2.13.0          nnet_7.3-20            
##  [46] Matrix_1.7-3            splines_4.5.0           tidyselect_1.2.1       
##  [49] effects_4.2-2           rstudioapi_0.17.1       abind_1.4-8            
##  [52] yaml_2.3.10             TMB_1.9.17              codetools_0.2-20       
##  [55] lattice_0.22-6          tibble_3.2.1            withr_3.0.2            
##  [58] flextable_0.9.8         askpass_1.2.1           evaluate_1.0.3         
##  [61] foreign_0.8-90          gridGraphics_0.5-1      survival_3.8-3         
##  [64] survey_4.4-2            zip_2.3.3               xml2_1.3.8             
##  [67] pillar_1.10.2           checkmate_2.3.2         reformulas_0.4.1       
##  [70] insight_1.3.0           plotly_4.10.4           generics_0.1.4         
##  [73] scales_1.4.0            minqa_1.2.8             xtable_1.8-4           
##  [76] glue_1.8.0              gdtools_0.4.2           Hmisc_5.2-3            
##  [79] lazyeval_0.2.2          tools_4.5.0             data.table_1.17.2      
##  [82] lme4_1.1-37             fs_1.6.6                mvtnorm_1.3-3          
##  [85] grid_4.5.0              mitools_2.4             tidyr_1.3.1            
##  [88] rbibutils_2.3           datawizard_1.1.0        colorspace_2.1-1       
##  [91] nlme_3.1-168            htmlTable_2.4.3         Formula_1.2-5          
##  [94] cli_3.6.5               rvg_0.3.5               textshaping_1.0.1      
##  [97] officer_0.6.9           fontBitstreamVera_0.1.1 viridisLite_0.4.2      
## [100] gtable_0.3.6            R.methodsS3_1.8.2       yulab.utils_0.2.0      
## [103] sass_0.4.10             digest_0.6.37           fontquiver_0.2.1       
## [106] ggplotify_0.1.2         TH.data_1.1-3           htmlwidgets_1.6.4      
## [109] farver_2.1.2            htmltools_0.5.8.1       R.oo_1.27.1            
## [112] lifecycle_1.0.4         httr_1.4.7              fontLiberation_0.1.0   
## [115] openssl_2.3.2           MASS_7.3-65
```
